# Condensation of the *Drosophila* Nerve Cord is Oscillatory and depends on Coordinated Mechanical Interactions

**DOI:** 10.1101/2021.02.24.432750

**Authors:** Katerina Karkali, Prabhat Tiwari, Anand Singh, Sham Tlili, Ignasi Jorba, Daniel Navajas, José J. Muñoz, Timothy E. Saunders, Enrique Martin-Blanco

## Abstract

During development, organs must form with precise shapes and sizes. Organ morphology is not always obtained through growth; a classic counterexample is condensation of the nervous system during *Drosophila* embryogenesis. The mechanics underlying such condensation remain poorly understood. Here, we combine *in toto* live-imaging, biophysical and genetic perturbations, and atomic force microscopy to characterize the condensation of the *Drosophila* ventral nerve cord (VNC) during embryonic development at both subcellular and tissue scales. This analysis reveals that condensation is not a unidirectional continuous process, but instead occurs through oscillatory contractions alternating from anterior and posterior ends. The VNC mechanical properties spatially and temporally vary during its condensation, and forces along its longitudinal axis are spatially heterogeneous, with larger ones exerted between neuromeres. We demonstrate that the process of VNC condensation is dependent on the coordinated mechanical activities of neurons and glia. Finally, we show that these outcomes are consistent with a viscoelastic model of condensation, which incorporates time delays due to the different time scales on which the mechanical processes act, and effective frictional interactions. In summary, we have defined the complex and progressive mechanics driving VNC condensation, providing insights into how a highly viscous tissue can autonomously change shape and size.

---

> “What utilitarian goal has nature pursued in forcing nervous system differentiation to these lengths?
>
> The refinement and enhancement of reflex activity, which protects the life of both the individual and the species” (Cajal, 1899).

## INTRODUCTION

Morphogenesis proceeds as a result of changes in cells proliferation, adhesion, differentiation and survival, and it is also the subject of mechanical inputs (Hogan, 1999) (Zhang and Labouesse, 2012) (Weber et al., 2011) (Heisenberg and Bellaïche, 2013). Further, during ontogenesis, all organs develop in synchrony to reach physiological optimization (Oliveira et al., 2014). In this scenario, how mechanics influences the final shape or size of an organ remains far from clear (Heisenberg and Bellaiche, 2013; LeGoff and Lecuit, 2015; Saunders and Ingham, 2019). A critical issue is that mechanical processes must be highly coordinated, while also accounting for geometric and scaling constraints (Amourda and Saunders, 2017).

Biological tissues display both elastic and viscous properties and are, in many cases, mechanically heterogeneous both in space and time (Serwane et al., 2017). They are constituted by active materials, and so standard equilibrium biophysical approaches are often insufficient to describe their behaviors. The material properties of tissues are thus key for the development of the organism (Mammoto and Ingber, 2010) (Miller and Davidson, 2013) (Mongera et al., 2018). However, understanding how the material properties of tissues impact the building and shaping of organs during development remains an open question.

Precise tissue organization is especially relevant when considering the functional complexity of the Central Nervous System (CNS) (Redies and Puelles, 2001). The complex architecture of the mature CNS is achieved through a well-known sequence of cellular events (Roig-Puiggros et al., 2020) (Tessier-Lavigne and Goodman, 1996). At the local level, tension forces contribute to the formation and maintenance of active synapses and the stabilization of neurites (Anava et al., 2009) (Kilinc, 2018). They also influence the shortening of neuronal processes, thus contributing to circuitry compactness (Franze, 2013). However, it is unknown which mechanical processes at the tissue-scale are involved in the spatial organization of neural architecture.

Here we fill this knowledge gap by determining how mechanical forces - generated at different scales - translate into tissue level sculpting of the entire *Drosophila* embryonic ventral nerve cord (VNC), as manifested by its pronounced shortening during condensation. The embryonic CNS is built stepwise by neuroblasts that delaminate from the embryonic neurectoderm in an invariant pattern, generating a diverse population of neurons and glia (Hartenstein and Wodarz, 2013). Neurons are unipolar and project their axons towards the neuropil. Cohesive axon bundles travel together and branch in the same or closely adjacent neuropil compartments creating stereotyped segmental structures (Landgraf et al., 2003) (Technau, 2008). Axon tracts include three longitudinal connectives that pioneer the neuropil of the VNC, and transverse pioneer commissures establishing contralateral connections (Lin et al., 1994). Neurons are supported by a complex scaffold of glia, which creates a meshwork of cortex processes required for stabilizing neurons’ positions (Beckervordersandforth et al., 2008). Only by integrating knowledge of the action of specific cell types along with long range mechanical forces can we begin to build a coherent model of tissue condensation.

Macroscopically, the VNC is formed along the extended germ band and exhibits a dramatic late shortening that further progresses in larval stages (Campos-Ortega and Hartenstein, 1985) (Page and Olofsson, 2008) (Olofsson and Page, 2005). It more than halves in length during the late embryonic stages. From an architectural viewpoint, the mechanisms modulating how the CNS gets shaped and how its composing elements are brought together into a mechanically stable functional structure are unknown.

To analyze the VNC condensation dynamics across multiple scales, we used four-dimensional confocal and light-sheet microscopy along with advanced image analysis. Velocity and strain maps revealed a complex morphogenetic kinematic, comprising alternate active and passive periods. Condensation during the active phases proceeds centripetally from both ends of the VNC and exhibits local oscillatory behavior. Further, spatial and temporal quantifications of material stiffness showed that the VNC displays a correlative, segmentally iterated, tensional landscape and stereotyped material stiffness inhomogeneities. To explain how these properties drive periodic oscillations, we built a viscoelastic model that revealed that they are consistent with having different viscous and elastic mechanical behaviors during tissue condensation. The combined experimental and theory results show that large-scale mechanical forces are essential for condensing and shaping the VNC. We assigned the ultimate acquisition of the VNC final shape and the generation of global force patterns to the concerted actions at a cellular scale of neurons and glia through the dynamic modulation of their cytoskeleton. Overall, this work reveals that the nervous system behaves as a solid viscoelastic tissue during condensation and that its biomechanical properties are key, in concurrence with a complex series of coordinated cellular actions, for its morphogenesis.

## RESULTS

### VNC condensation dynamics

To understand the mechanics underlying condensation at the global tissue scale we characterized its progression *in vivo*, from the initiation of germ band retraction to larval hatching. Midway through embryogenesis, the *Drosophila* VNC undergoes dramatic condensation along the AP axis, shortening from over 700 to less than 250 µm (**Figure S1** and **Movie S1**). This process depends on different cellular events: the remodeling of the extracellular matrix (ECM) by hemocytes; the cytoskeletal dynamics of glia and neurons; and regulated apoptosis (Olofsson and Page, 2005) (Page and Olofsson, 2008) (Evans et al., 2010). We live-imaged Fasciclin 2 (Fas2)-GFP embryos (Buszczak et al., 2007) by confocal microscopy and embryos expressing the nuclear marker Histone2A-mCherry by light-sheet microscopy (Krzic et al., 2012). To reconstruct the VNC three-dimensional morphology from stage 16 onwards we had to overcome the movements undertaken by embryos. We employed an image processing pipeline that “detwitched” embryos by digitally re-locating the VNC along the central midline at each time-point from *in toto* light-sheet images **(Figure 1A, Movie S2** and Experimental Procedures).

**Figure 1:**
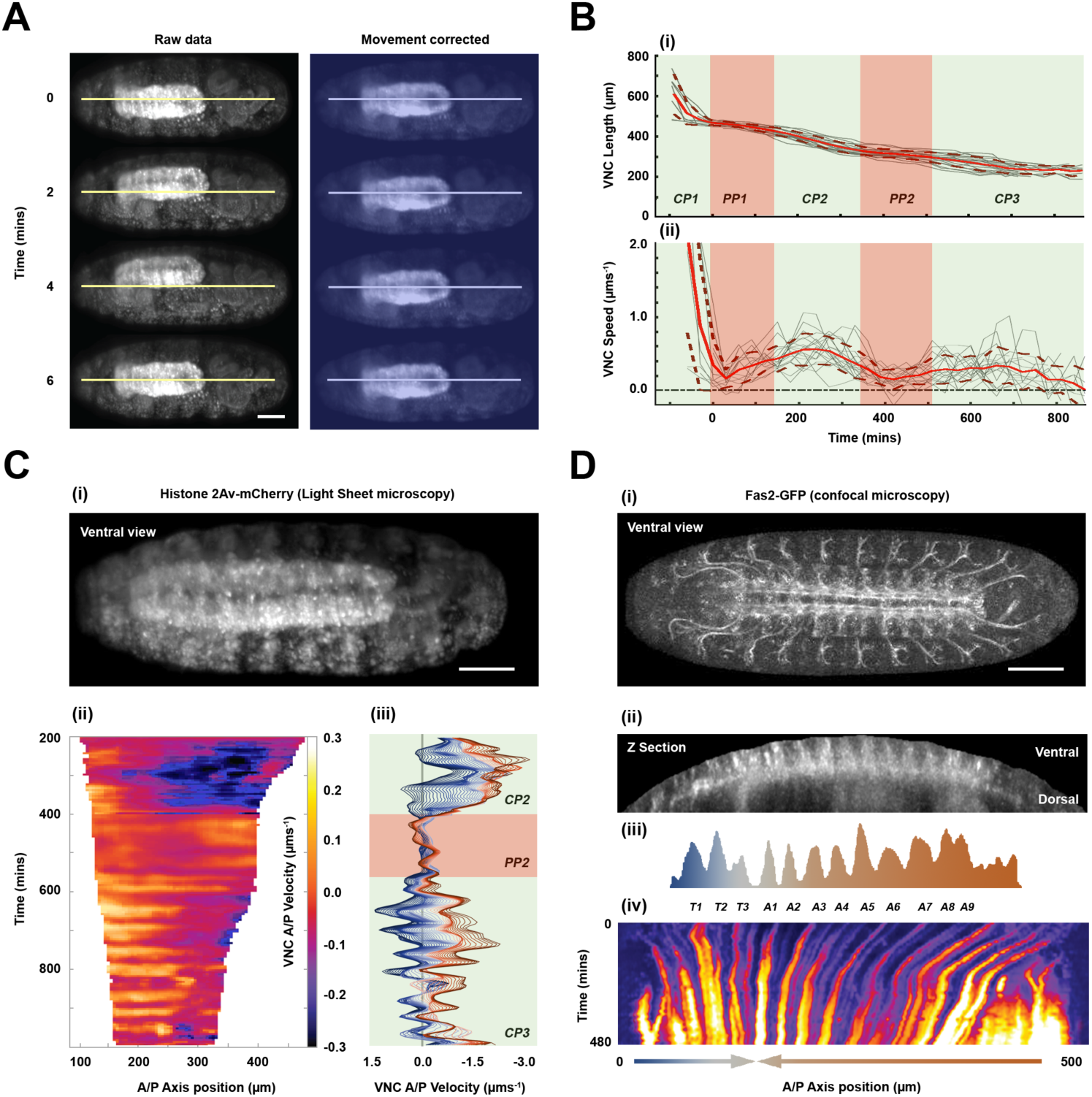
Dynamics of VNC condensation. **A)** Snapshots at 2-minute intervals from a time lapse **(Movie S2)** recorded by multi-view light-sheet imaging of a live Histone 2Av-mCherry embryo (ventral view – late stage 17). mCherry labeling marks all nuclei and was used to correct the embryo twitching (see Experimental Procedures); raw data is shown on the left and “detwitched” images (blue masked) on the right. In all images, anterior is to the left and posterior to the right. Lines indicate the ventral midline. Scale bar 50 µm. **B)** Quantification of VNC length (**i**) and condensation speed (**ii**) as function of time. Condensation (CP1, CP2 and CP3) and pause (PP1 and PP2) phases are masked in pale green and red respectively. As a convention for this and all subsequent figures (unless stated otherwise), t=0 corresponds to the onset of the VNC pause phase (PP1), at the end of germ band retraction. Means (solid) and SD (dashed) are represented by red lines. Gray lines represent individual embryos (n=11 embryos). **C)** Condensation speed spatiotemporal dynamics. (**i**) Snapshot of a live Histone 2Av-mCherry embryo monitored by light-sheet imaging at stage 16. Scale bar 50 µm. (**ii**) Velocity kymograph derived from PIV analysis (Experimental Procedures) along the VNC. For this and all subsequent figures, position=0 along the AP axis corresponds to the hinge between the brain lobes and the VNC proper. Time axis (top to bottom) was defined as in **(B)**. Color-coded positive and negative values of velocity correspond to posterior-ward and anterior-ward displacements respectively. (**iii**) Representation of velocity profiles along the whole condensation process (CP2, PP2 and CP3), with 5-minute resolution, for all points along the AP axis from the most anterior (darkest blue) to the most posterior (darkest red lines) VNC positions. **D)** Kymograph along the VNC length from a live embryo expressing Fas2-GFP. (**i**) Ventral view from a confocal microscopy acquisition **(Movie S3)**, at stage 16. Scale bar 50 µm. (**ii**) Stage 16 embryonic VNC, re-sliced over the Z-axis. (**iii**) Fluorescence intensity peaks mark individual segments landmarks (color coded as in **(C)**). Time and AP axis positions are as in **(B)** and **(C)**. (**iv**) Kymograph of condensation, with arrows denoting condensation direction.

We generated length and velocity profiles for the VNC throughout condensation **(Figure 1B)**. These revealed that condensation proceeds in five dynamic steps. First, the VNC pulls back in association with the retraction of the germ band until its posterior end positions near the posterior tip of the embryo (compaction phase 1 – CP1). The condensation speed in this phase follows that of the germ band, which is variable at first and sustained later (Lynch et al., 2013). Second, the VNC reaches an almost stationary phase by the end of germ band retraction (end of stage 13). This phase lasts for around two hours at 25°C, up to the end of dorsal closure and head involution by late stage 15 (pausing phase 1 – PP1). Third, the VNC, uncoupled from the epidermis and mesoderm, rapidly initiates contraction (compaction phase 2 – CP2). Fourth, condensation reaches a second resting period of around two hours (pausing phase 2 – PP2). Fifth and last, the VNC undergoes a final slow progressive compaction, concurrent with peristaltic embryo movements, up to the end of stage 17 (compaction phase 3 – CP3). Variability in VNC length between embryos along the whole process is very small (<10%) except during the germ band retraction-associated CP1, indicating that VNC condensation is a robust process structured in alternating active and passive phases **(Figure 1B)**.

### VNC condensation is oscillatory

Initially, condensation of the emerging VNC passively follows the movements of the germ band. The successive autonomous phases of contraction are, on the other hand, active processes. We undertook a comprehensive analysis of these later steps by quantifying, using particle image velocimetry (PIV) (Vig et al., 2016), the local velocities along the whole length of the VNC as it condenses (phases CP2, PP2 and CP3) **(Figure 1C**, **Movie S2)**. Remarkably, we found that, after head involution, condensation is oscillatory, both during the CP2 and CP3 phases, with contractile periods on a time scale of around 30 minutes. The frequency of the oscillations is quite regular, while their amplitude is temporally varying. Contractile oscillations are observed at the anterior and, most prominently, at the posterior of the VNC with opposing directionalities **(Figure 1C)**. They lead to the bidirectional convergence of the VNC towards a central stationary domain located between the third thoracic and the first abdominal segments (**Figure 1D** and **Movie S3**).

In summary, the late active shortening periods of VNC condensation are not monotonic. Tissue-scale oscillations suggest a sophisticated spatiotemporal mechanical coordination across the whole tissue.

### Material properties of the VNC vary both spatially and temporally during development

The unexpected complex kinematics of condensation hints to potential spatiotemporal changes in VNC material properties. To characterize VNC stiffness during condensation across time and in different parts of the tissue we used Atomic Force Microscopy (AFM). The elastic Young’s modulus (E), a stiffness proxy, was measured segment-to-segment at the midline and at the lateral neuropile of stage 14 and late stage 16 embryos (**Figure 2A-C** and Experimental Procedures).

**Figure 2:**
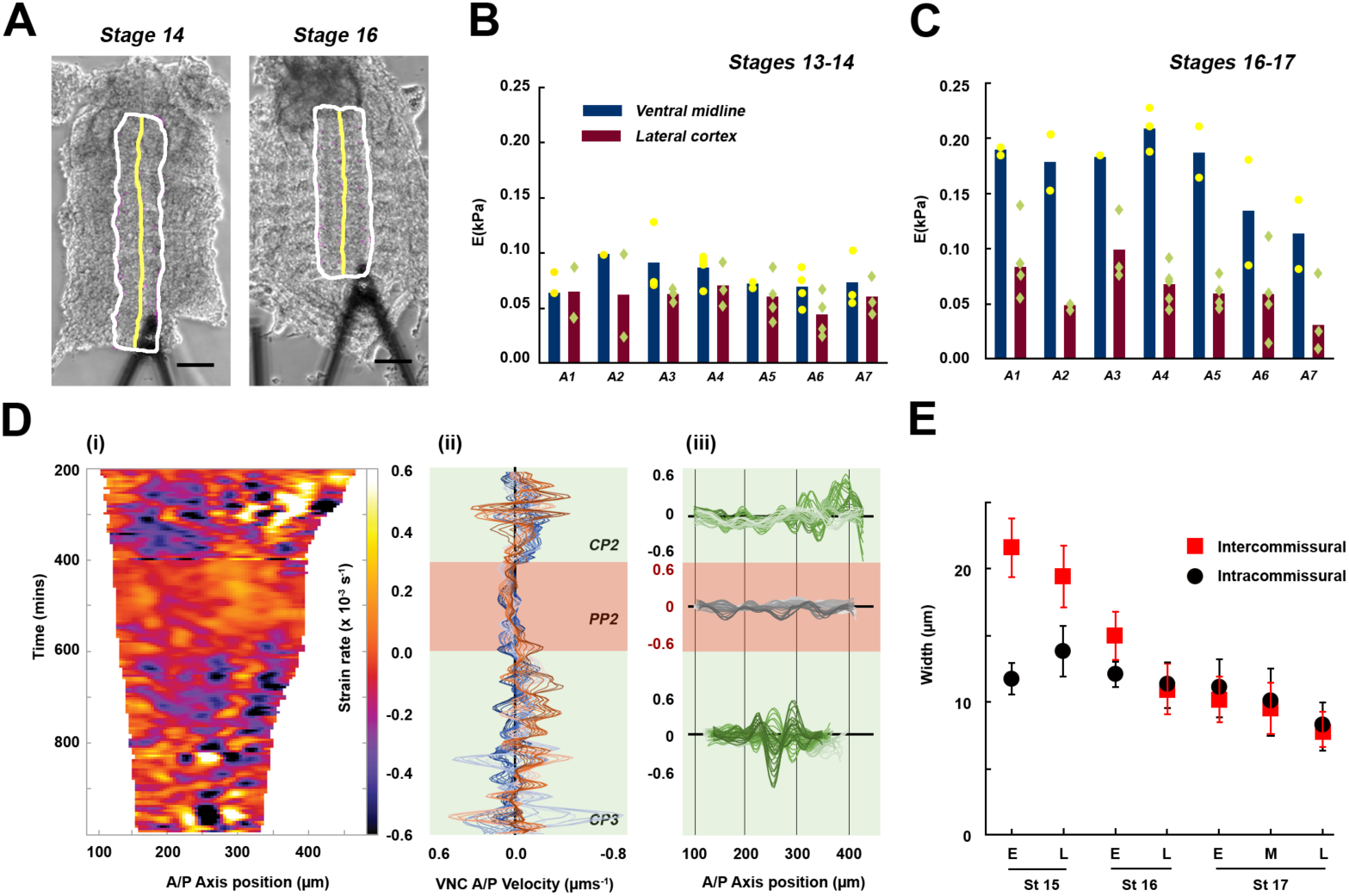
Characterization of the VNC material properties and local dynamics along its condensation. **A)** Representative images of flat dissected embryos in stages 14 and stages 16, respectively. VNC perimeter (white line) and midline (yellow line) are highlighted, with the AFM cantilever head shown. Anterior is to the top. Scale bar 50 µm. **B)** Measured tissue stiffness (E) for dissected VNC at early stages (13-14). Bars denote mean values at each abdominal segment A1 to A7. Mean tissue stiffness was measured at the midline (blue) and at lateral positions of the cortex (red). Dots and diamonds correspond to individual measurements. **C)** as **(B)** but for later, stage 16-17, samples. **D)** (**i**) Kymograph of VNC strain rates, from data in Figure 1C (see Experimental Procedures). (**ii**) Representation of strain rates profiles during condensation (CP2, PP2 and CP3), with 5-minute resolution, from the most anterior (darkest blue line) to the most posterior (darkest red line) VNC positions. (**iii**) Distribution of strains in VNC along the AP-axis for all time points (earliest light to latest dark lines) during the phases CP2 (green), PP2 (gray) and CP3 (green). **E)** Average size (and SD) of intra- and inter-commissural domains from early (E) and late (L) stage 15, early (E) and late (L) stage 16 and early (E), middle (M) and late (L) stage 17 embryos as the VNC condenses. Data was collected from 7-10 measurements per time point from two embryos.

At stage 14 of embryogenesis (PP1 phase), before the VNC active contractions initiate, E was 0.08 ± 0.01 KPa (mean ± SD, n=15) in the midline, and 0.06 ± 0.01 KPa in the lateral regions (abdominal segments 1 to 5) **(Figure 2B)**. No statistical differences were found either between midline and lateral positions or along the AP axis (p > 0.05). At late stage 16 (late CP2), the midline stiffness increased significantly in comparison to the lateral cortex area to stage 14 (p < 10^-3^). E in the midline was 0.17 ± 0.03 KPa and 0.06 ± 0.02 KPa (n=15) in the lateral regions. We also found that stiffness varied along the AP axis (**Figure 2C**); the anterior region was stiffer, with a dramatic decrease towards the most posterior segments (see also **Figure S2**).

Therefore, consistent with previous measurements in neural tissues from different organisms or from individual neurons (Spedden and Staii, 2013), the *Drosophila* embryonic VNC is extremely soft. Our results indicate that the neural tissue stiffens with time in an axially graded fashion. The central domain of the embryonic VNC, where most axons bundle, becomes more rigid than the lateral domains, where the somata are predominantly located.

### The tensional landscape of the VNC is temporally and spatially patterned

To infer large tissue scale forces, we determined local strain rates. Strain rates measure how rapidly neighboring regions move relative to each other within a given domain (Petridou and Heisenberg, 2019). They are related to the resulting tissue stresses, which also depend on the viscoelastic tissue properties; its bulk viscosity and the local shear rate (Experimental Procedures). The strain rate maps reveal that tissue deformations get restricted to specific subdomains as the VNC condenses (**Figure 2D** and **Movie S2**). At the active CP2 and CP3 phases, the strain rates display alternating positive and negative values along the AP axis. The strain rate has a distinct change in magnitude immediately after the second pause phase, before slowly tending back towards zero. Distinct strain rate domains appear to correspond to discrete contractile regions iteratively repeated along the VNC. Direct measurements of the distances in between inter- and intra-commissural domains show that the contractile regions map to the space between the posterior commissure of one neuromere and the anterior commissure of the next (intercommissural) **(Figure 2E)**.

Together, the strain rates maps and AFM data indicate that the VNC is mechanically organized in repeating units, and its mechanical landscape is temporally modulated. To confirm these propositions, we studied the response to mechanical perturbation of the VNC during condensation. We utilized laser microsurgery to sever the VNC at specific times and positions (**Figure 3A**, **Figure S3A** and Experimental Procedures). Cutting transversally to the AP axis in the intercommisural domains between the abdominal neuromeres of stage 11-14 embryos resulted in an isotropic recoil faster than in intracommissural (neuromere) regions (**Figure 3B**, **Figure S3B** and **Movie S4**). Our data suggest that the intercomissural domains are under significantly higher tension than the intracommissural during the stationary phase (PP1). By contrast, during the late condensation phase (CP2/PP2 - stages 16-17), tissue recoil after cutting was essentially lost **(Figure 3C)**. Importantly, severing an individual intercommissural space does not affect adjacent neuromeres that continue to condense (**Figure 3D** and **Movie S5**); they act as apparently independent units.

**Figure 3:**
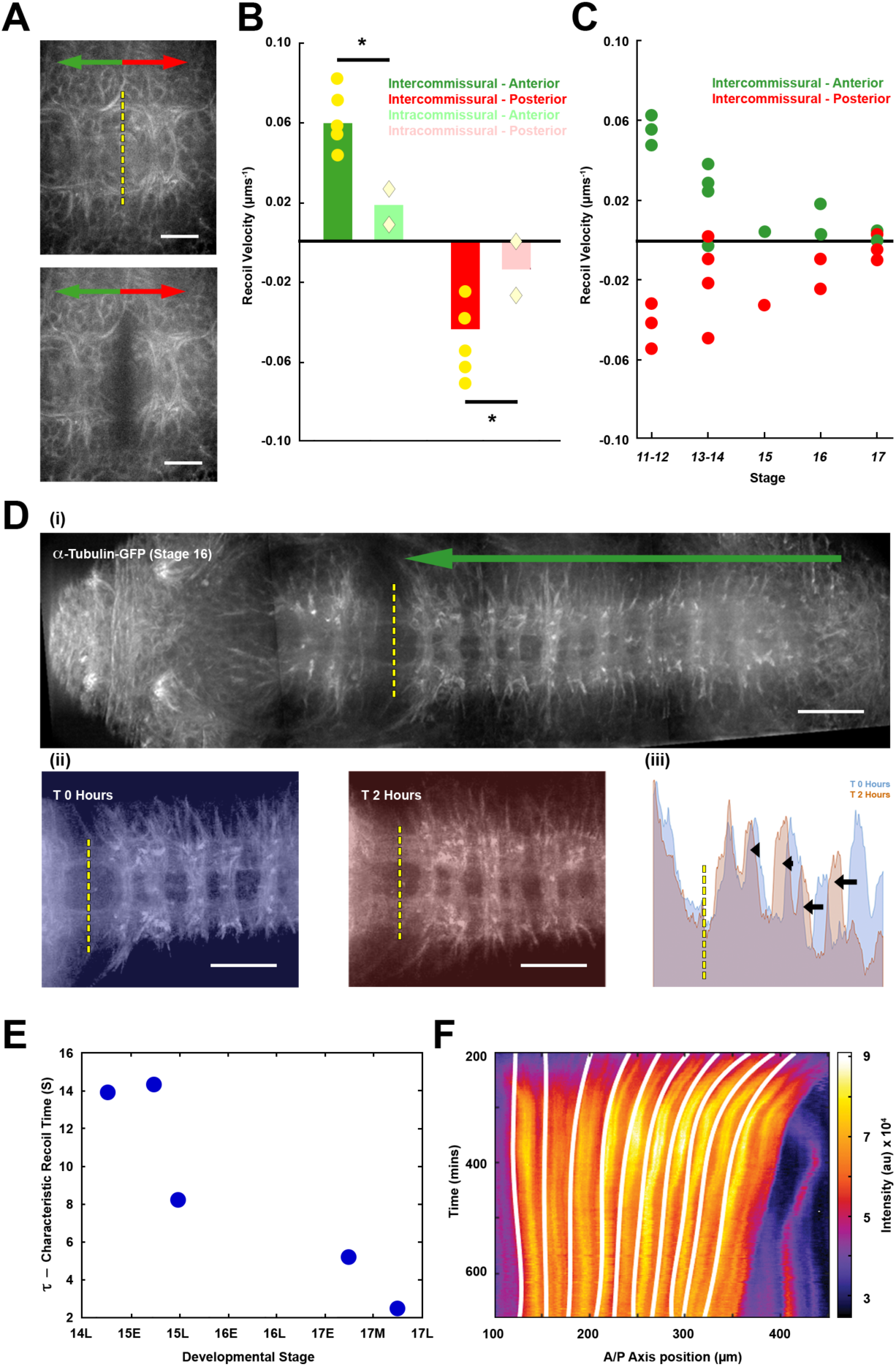
VNC response to laser microsurgery during condensation and tissue tension. **A)** Representative images of stage 14 embryos, expressing alpha Tubulin-GFP, before and after laser ablation. The yellow dashed line highlights the position of the laser cut (intercommissural), while green (anterior) and red (posterior) arrows indicate tissue recoil directionality **(Movie S4)**. Scale bar 10 µm. **B)** Tissue recoil velocity after ablation at intercommissural (dark) and intracommissural (pale) domains, on stage 14 embryos. Bars represent mean recoil velocity of anteriorly (green) and posteriorly (red) retracting tissue. Individual measurements are denoted by yellow dots and diamonds. * p < 0.05. **C)** Recoil velocity of anteriorly (green) and posteriorly (red) retracting domains after VNC ablation at different stages of embryonic development (n=12 embryos). **D)** (**i**) tiled image of a stage 16 embryo expressing alpha Tubulin-GFP after laser cutting the intercommissural domain between abdominal segments A1 and A2. The green arrow marks the direction of tissue condensation. (**ii**) Snapshots, immediately post-ablation (masked blue), and 2 hours later (masked red), from **Movie S5**. Scale bar 20 µm. (**iii**) Superimposed fluorescence intensity profiles of both time points. Black arrows indicate the magnitude of the anterior-ward displacement of individual segmental landmarks over the analyzed period. **E)** Characteristic recoil time τ at different embryonic stages computed from the rate of recoil after laser ablation at the intercommissural domain. **F)** Kymograph of the VNC during condensation (Fas2-GFP expressing embryo). White curves correspond to fourth order polynomial fitting of the points of maximum compression as deduced from the viscoelastic FE model of the VNC (Experimental Procedures). (See also **Figure S3E** and **Movie S6**).

Laser cuts also enable an approximate characterization of the viscoelastic properties of the VNC by analyzing the recoil rate (see Experimental Procedures). Though we cannot discount possible viscous differences between the inter- and intracommissural domains, such potential variations are likely negligible as their tissue composition is equivalent. By severing the intercommissural domains at different time points we found a strong reduction of contractility and viscosity as the VNC condenses **(Figure 3E)**.

To evaluate the stress evolution along the VNC during condensation, we constructed a three-dimensional Finite Element model (FE). We mapped the measured velocity field onto the FE model and reconstructed the strain and stress fields along the VNC (**Figure 3F**, **Figure S3C-D**, **Movie S6** and Experimental Procedures). The evolution of the stress profiles along the AP-axis and the superposition of the stress minima (compression) onto the phase contrast kymographs, confirmed that maximum compression is sustained at the intracommissural domains. Further, the active stress increased over time in the intercommissures during the autonomous condensation phase **(Figure S3E)**. This observation points to a potential scenario in which the distribution of tension reflects the spaced contractions of the tissue (segments contract as units), and subsequently the progression of condensation.

### Oscillations are an emergent property of a viscoelastic tissue

The contraction of the intercommissural domains in between neuromeres explains the condensation of the VNC, but how do they coordinate? Can this coordination explain the origin of the global oscillations? To tackle these problems, we developed a one-dimensional rheological model that incorporates the essential viscoelastic properties of the VNC along with a delayed active contractility. At a particular time, *t*, the VNC is taken to have a rest length, *L(t)*. This internal variable depends on time, as the system gradually condenses. The actual length of the VNC at a particular time, *l(t)*, can differ from *L(t)* due to tissue relaxation, active reorganization and stress acting on the system. We define Δ*L=l(t)-L(t)*, representing the difference in VNC length from its rest length at time *t*. Using this, we can represent the change in the rest length as a function of time by

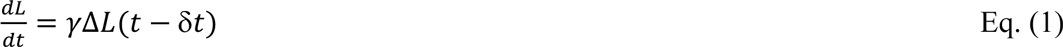

where *γ* is the remodelling rate, which measures the rate at which the tissue adapts its rest-length. The time delay, δ*t* in Eq. (1) represents the delay between the current strain measure Δ*L* and the active remodeling of the VNC through its rest-length *L*. The VNC condenses within the densely packed environment of the embryo, it is surrounded by the neural lamella (Meyer et al., 2014) and it is connected early on to the underlying epithelia and later by intersegmental and segmental nerves to the developing muscles and peripheral sensory organs. We also incorporate potential effects of surrounding tissues in the model. We added to the rheology a frictional term proportional to the apparent VNC length rate *dl/dt,*

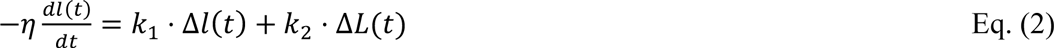

where *η* is the friction coefficient, *k*_1_ is the purely elastic component of the VNC, Δ*l*(*t*) = *l*(*t*) − *l*_0_(*t*) (where *l_0_* is the characteristic elastic length scale), and *k*_2_ represents the stiffness of the viscoelastic component of the VNC, with a dynamic rest-length *L(t)*. (**Figure 4A**). The combination of Eqs. (1) and (2) yields a viscoelastic model with a delayed viscous response, which has the ability to exhibit oscillatory behavior (Muñoz et al., 2018).

**Figure 4:**
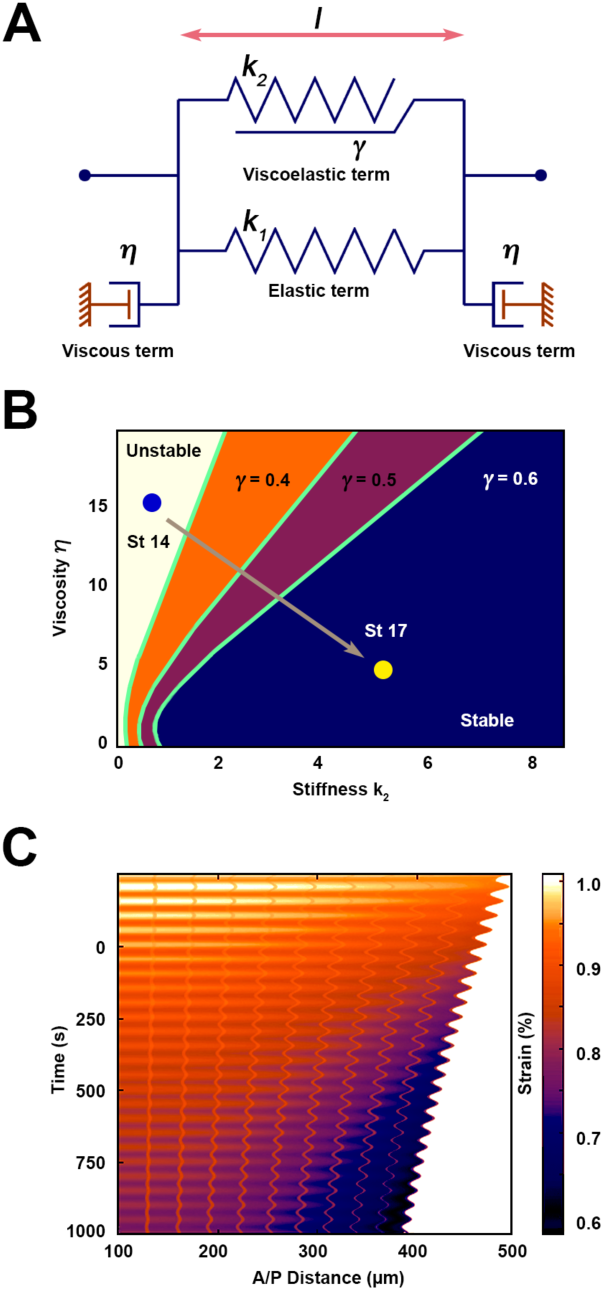
Rheological model of VNC condensation. **A)** Scheme of one-dimensional rheological model. A viscoelastic term with variable rest-length *l* has stiffness *k*_2_ and remodeling rate γ (see Eq. (1) in Results). The VNC is taken to have an elastic component in parallel, with stiffness *k*_1_. The model also includes viscous contact to the external environment, denoted by η. **B)** Phase diagram in the parameter space *k*_2_ - η, showing that reduction of η and increase of *k*_2_ stabilizes the oscillatory behavior. Points St 14 and St 17 represent material values and transition from early to later stages of VNC development, with a stabilizing effect. **C)** Kymograph of numerical simulation showing the oscillatory behavior of strains as a function of time. Simulations with other parameter values showing unstable responses are shown in **Figure S4**.

Similar rest-length models have recently been used in the context of embryogenesis (Cavanaugh et al., 2020) (Doubrovinski et al., 2017) (Sumi et al., 2018), epithelia remodeling (Clement et al., 2017) (Staddon et al., 2019) and stress relaxation of monolayers (Khalilgharibi et al., 2019). However, the stability of the model with the delay rheology in Eq. (1) has only been analyzed in the absence of environmental viscous effects (Muñoz et al., 2018). Eqs. (1)-(2) form a system of Delay Differential Equations that can be analyzed through their characteristic equation (Stépán, 1989) (Erneux, 2009) or numerically. Depending on the parameters η, γ, δt, *k*_1_ or *k*_2_, the apparent length *l*(*t*) can exhibit a stable regime (with no oscillations or oscillations showing a diminishing amplitude) or unstable oscillations (with increasing amplitude). The phase diagram in **Figure 4B** shows that decreasing values of *k*_2_ and γ render the system unstable, while decreasing values of viscosity η render the oscillations more stable. These results are consistent with the eventual stabilization of the VNC as its stiffness increases **(Figure 2B-C)** and its viscosity is progressively reduced **(Figure 3E)**. The kymograph in **Figure 4C** shows an example of the stable oscillatory regime (see **Figure S4** for other scenarios). Overall, our reduced one-dimensional model can explain the emergence of the periodic contractions as a consequence of time delays induced from the material properties of the VNC and possible effective friction between the neural cortex and the surface glia. As the VNC stiffens during development, these oscillations are stabilized, ensuring robust formation of the VNC.

### VNC condensation requires significant mechanical contribution from glia

Can differences in material properties and emergent oscillations be connected to cell behaviors? Previous analyses of VNC condensation have defined differing roles for glial cells in the removal of dead cells, or the constriction itself (Shklyar et al., 2014) (Olofsson and Page, 2005). Considering the complex mechanics of VNC condensation, we next asked if neurons or glia – through their cellular scale actions - play a mechanically active part in modulating tissue scale behavior and, if they do, we aimed to determine their effects on the VNC material properties.

To approach these questions, we first genetically ablated either neurons or glia by overexpressing the proapoptotic gene Grim (Chen et al., 1996) employing the pan-neural Elav-Gal4 and the glia Repo-Gal4 drivers **(Figure 5A-D)**. Excessive neuronal cell death heavily distorted the organization of the axonal scaffold **(Figure 5A)**, and, to a lesser extent, the VNC condensation **(Figure S5A)**. Though excessive glia apoptosis was not observed in this condition, glial cells mispositioned, probably in response to steric constrains resulting from alterations in axonal morphology **(Figure 5B)**. Expressing Grim in glia promoted, indirectly, slight alterations in the 3D axonal scaffold **(Figure 5C)**. Most of the glia were removed **(Figure 5D)** and resulted in a strong failure of the condensation process **(Figure S5A)**. Loss of glia also altered the VNC shape, which was distorted in a stereotyped way (**Figure 5E** and **Movie S7**). Employing light sheet microscopy (**Figure S5B** and **Movie S8**) we observed that in the absence of glia, VNC condensation arrests at the last contraction phase (CP3) **(Figure 5F)**. PIV analysis of VNC condensation revealed a weakly structured segmented pattern, with a substantial reduction of VNC strain rates at all time points and positions, and the loss of contractile oscillations **(Figure S5C)**.

**Figure 5:**
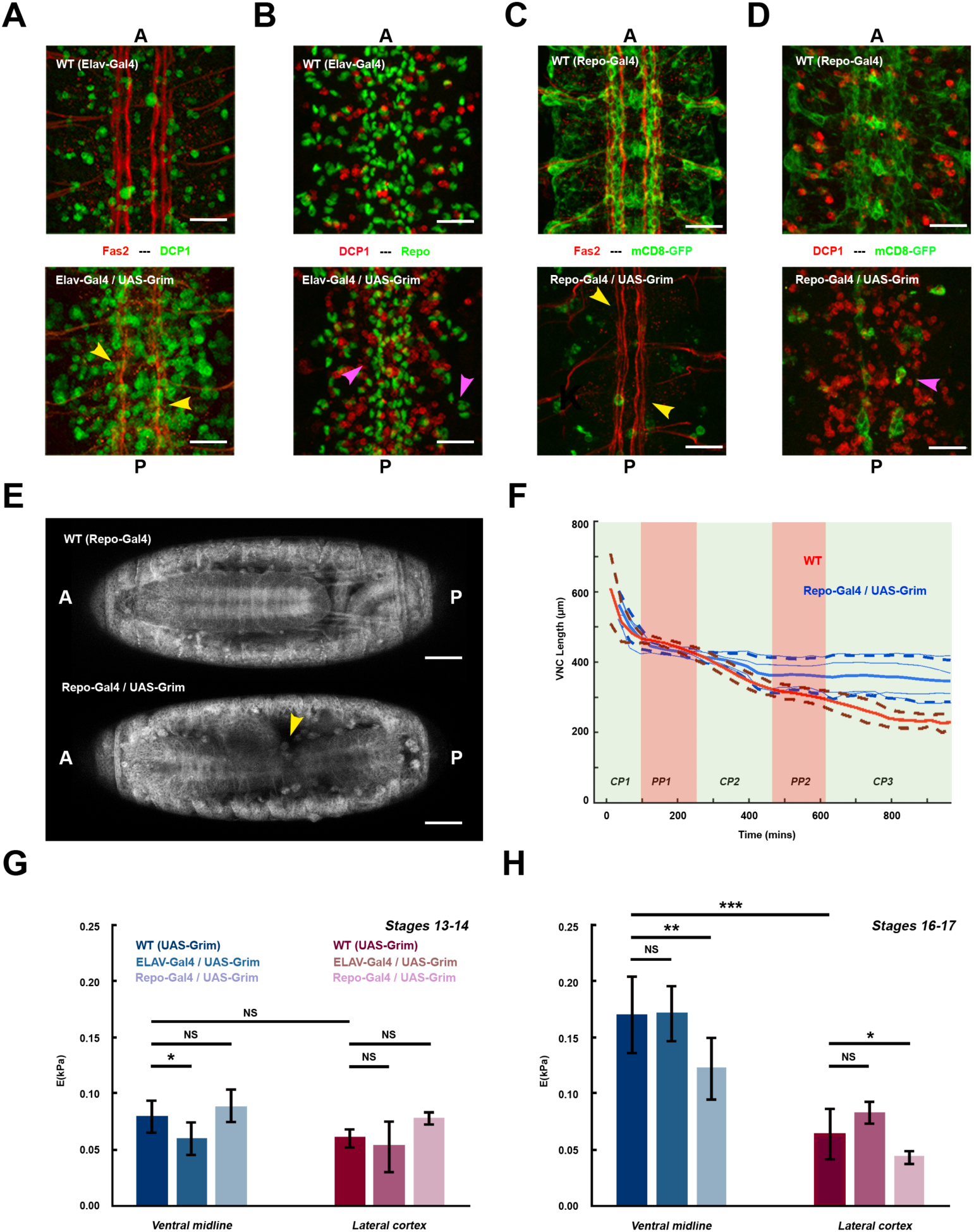
Both neurons and glia participate in the architectural organization of the VNC and its condensation. **A)** CNS Flat-preps of WT (top) and Elav-Gal4>UAS-Grim (bottom) embryos, at stage 16, immunostained for Fas2 (red) and Dcp1 (green). Yellow arrowheads point to the disrupted axonal network. **B)** Embryos of the same genotype as in **(A)**, immunostained for Dcp1 (red) and Repo (green). Pink arrowheads point to misplaced glia. **C)** CNS Flat-preps of WT (top) and Repo-Gal4:UAS-mCD8-GFP>UAS-Grim (bottom) embryos, at stage 16, immunostained for Fas2 (red) and GFP (green). Yellow arrowheads point to the disrupted axonal network. **D)** Embryos of the same genotype as in **(C)**, immunostained for Dcp1 (red) and GFP (green). Pink arrowhead points to surviving glia. Scale bar 10 µm in **A-D**. **E)** Snapshots from time lapse recordings of WT (Top) and Repo-Gal4>UAS-Grim (bottom) embryos, in an alpha Tubulin-GFP background (ventral view –stage 17) **(Movie S7)**. Yellow arrowhead points to the VNC misshaped buckling. Scale bar 50 µm. AP axis orientation is indicated. **F)** Quantification of VNC length as a function of developmental time in WT (red, n=11) and Repo-Gal4>UAS-Grim (blue, n=4) embryos marked with *elav*:mCD8-GFP by confocal imaging. Solid and dashed lines show mean and SD values respectively. **G)** Tissue stiffness (E) measured by AFM for dissected VNCs at early stages (13-14), from WT, Elav-Gal4>UAS-Grim and Repo-Gal4>UAS-Grim embryos. Bars denote mean values at the ventral midline (blue) and at lateral cortex areas (red). *p < 0.05, **p < 10^-2^ and ***p < 10^-3^. **H)** As **(G)** but for later stage 16-17 embryos.

We explored by AFM whether neurons or glia modulate the VNC material properties. AFM measurements were performed at the early stationary PP1 (stage 14, **Figure 5G**) and the second active condensation CP2 phases (stage 16, **Figure 5H**) after inducing targeted cell death. Before active contraction, VNC rigidity slightly decreased at the midline after neuron ablation but it was not affected by glia depletion. On the contrary, during the active CP2 phase, significant softening upon glia removal was observed, both at the midline and at lateral positions, while neuronal ablation had no effect (**Figure 5H**).

In summary, both glia and neurons contribute to the active contraction of the VNC and modulate its material properties. However, while glia have a major contribution on both of these aspects of the condensation process, the impact of neurons appears to be more subtle; they just influence the structural organization of the neuropile and not VNC material properties.

### Myosin-mediated contractility in neurons and glia is required for VNC condensation

VNC condensation is an active process demanding mechanical efforts by both neurons and glia. Yet, how is intracellular force generated within these cells and how does this translate to tissue scale effects? To evaluate the mechanical impact that the active cytoskeleton may have in condensation, we analyzed actomyosin contractility in both cell types.

Pan-neural expression of a Zipper (Zip) RNAi transgene (Non-Muscle Myosin II heavy chain) resulted in major defects in the structural scaffold of the VNC and in VNC condensation failure (**Figure 6A** and **Figure S6A**). No increase of cell death was observed. Likewise, interference with Zip expression in glia also resulted in condensation defects **(Figure S6A)**. Glia looked smaller and failed to migrate properly. Neuronal longitudinal axonal tracts are found closer towards the midline **(Figure 6B)**. Analyzing VNC condensation dynamics, we found that Zip knockdown led to distinct spatiotemporal alterations (**Figure 6C-E** and **Movie S9**). Abolishing neuronal contractility by depleting Zip in neurons blocks VNC condensation by affecting all phases from CP2 onwards **(Figure 6D)**. Interestingly, although no condensation progression was detected, segmentally iterated strain rate differences were present. On the contrary, depletion of Zip in glia resulted in cessation of condensation at the PP2 phase **(Figure 6E)**. Last, FE model analyses indicate that upon blocking contractility in glia, no apparent compression takes place in intercommissural regions, and the strain rate pattern fades away (**Figure 7A-B** and **Movies S10** and **S11**).

**Figure 6:**
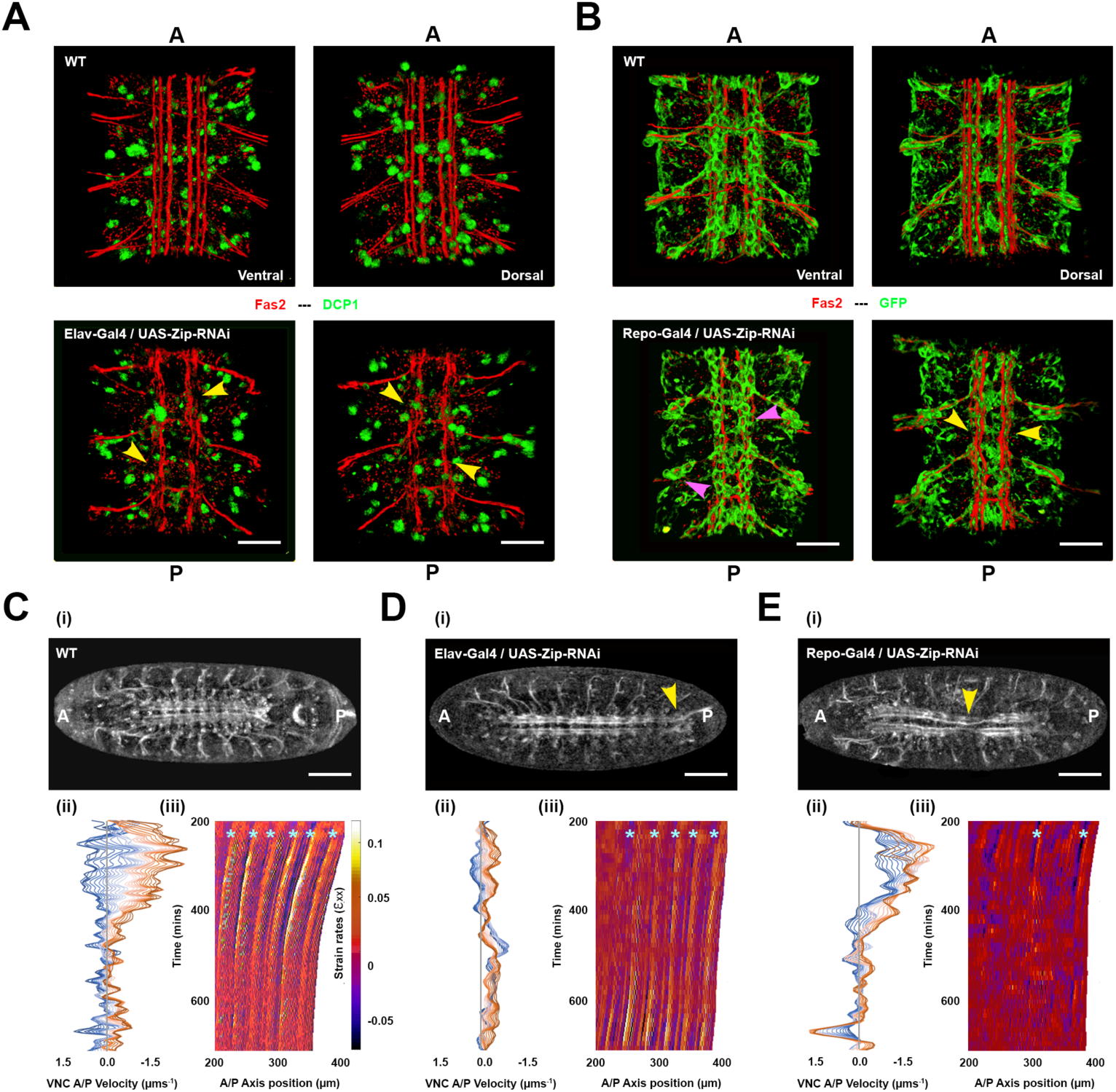
Active contractility in neurons and glia have distinct roles for VNC architecture and condensation. **A)** Ventral and Dorsal 3D views of dissected, stage 16, WT (top) and Elav-Gal4>UAS-Zip-RNAi (bottom) embryos, immunostained for Fas2 (red) and Dcp1 (green). Yellow arrowheads point to the disrupted axonal network. **B)** Ventral and Dorsal 3D views, equivalent to (**A**), of stage 16, WT (top) and Repo-Gal4:UAS-mCD8-GFP>UAS-Zip-RNAi (bottom) embryos, immunostained for Fas2 (red) and GFP (green). Pink arrowheads point to misplaced glia. Yellow arrowheads point to the disrupted axonal network. **A-B** Scale bar 10 µm. **C-E)** Condensation dynamics in control **(C)**, Elav-Gal4>UAS-Zip-RNAi **(D)** and Repo-Gal4>UAS-Zip-RNAi **(E)** embryos **(Movie S9)**. (**i**) Snapshots of live embryos, expressing Fas2-GFP, monitored by confocal imaging, at stage 17. Yellow arrowheads point to the posterior tip of the uncondensed VNC **(D)** and to the VNC misshaped buckling (**E**). Anterior is to the left. Scale bar 50 µm. (**ii**) Representation of velocity profiles during condensation along the AP axis, from the most anterior (darkest blue) to the most posterior (darkest red line) VNC positions (as in Figure 1C). (**iii**) Kymograph of strain rates along the VNC (as Figure 2D). Cyan marks point to strain oscillations, which are strongly diminished upon reduction of glia contractility.

**Figure 7:**
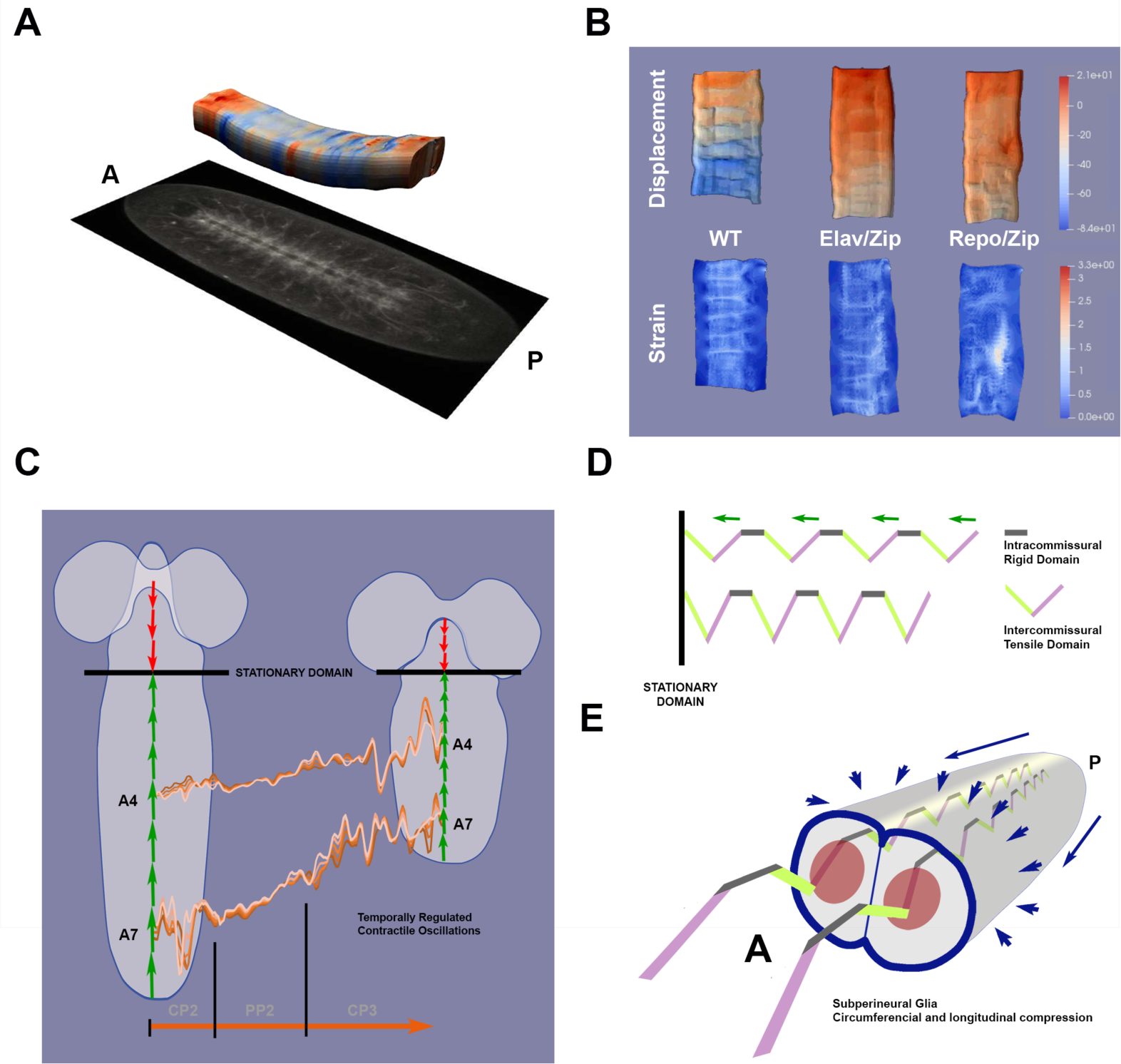
Neurons and Glia cooperate to lead the oscillatory character of VNC condensation. **A)** Snapshot from the 3D representation **(Movie S10)** of the segmental 2D strain pattern of the VNC during condensation, in WT animals. The actual 3D meshwork (top) is aligned to the corresponding raw image (bottom). AP axis orientation is indicated. **B)** Displacements and strains along the VNC, in WT embryos and in embryos with pan-neural (Elav-Gal4>UAS-Zip-RNAi) or pan-glial (Repo-Gal4>UAS-Zip-RNAi) non muscle Myosin II knockdown, at equivalent developmental times. (Snapshots from **Movie S11)**. **C)** Cartoon summarizing the VNC condensation oscillatory regime during the CP2 and CP3 stages (examples at the level of the abdominal segments A4 and A7, data from **Movie S2** – see Figure 1C), as well as pointing to the opposing displacements of the thoracic (red) and abdominal (green) segments towards the central stationary domain. **D)** Cartoon presenting the segmentally iterated intercommissural and intracommissural domains of the axonal network before (top) and after (bottom) condensation. Their mechanical properties (rigid or tensile) are shown. This representation depicts the first three abdominal segments actively contracting (green arrows) towards the thorax/abdomen stationary domain. **E)** Cartoon presenting in 3D the VNC internal segmentally iterated axonal network (described in **D**) surrounded by the glial shell (Subperineural Glia) with centripetal and longitudinal contractile capability (blue arrows). AP axis orientation is indicated.

These data support a model in which neuronal contractile capability from within the VNC (neurons) is permissive for its compaction at all stages but it is not sufficient. The glia exerts an external compressive force at the VNC surface, spatiotemporally regulated, that exploits the segmentally iterated architectural organization of the neuronal network to accommodate the periodic tensional pattern of the VNC into oscillatory condensation events **(Figure 7C-E)**.

## DISCUSSION

Our characterization of the spatiotemporal progression of VNC condensation provides essential cues for understanding the sequential steps involved in the acquisition of the final shape of the embryonic CNS. In discussing the spatial design of the nervous system we have to consider some specific constraints: (1) first, and foremost, the organism symmetry (bilateral or radial); (2) the spatial organization of its locomotor and sensorial machinery; and (3) the need to create an integrated and connected system that links the organization of muscles and sensory organs so that impulses can be centrally controlled and coordinated (Bullmore and Sporns, 2012; Swanson, 2007). The condensation of the VNC, within the global CNS developmental plan, must satisfy these constraints. During condensation, the VNC maintains its iterated axonal connections to all segments’ muscles and sensory organs.

### The condensation of the VNC is oscillatory and serves specific morphogenetic purposes

Condensation is a common morphogenetic event (Hall and Miyake, 2000), affecting the compaction of mesenchymal tissues. Condensation plays an important role at the earliest stages of organogenesis (*e.g.* cartilage, bone, muscle and tendon) (DeLise et al., 2000) and in shaping neural ganglia, both in arthropods (Bullock and Horridge, 1965) and vertebrates (Stark et al., 1997). In most condensation cases, cells get together by migratory accretion and the final form of the tissue is acquired by sustained anisotropic cell supply or intercalary growth (Christley et al., 2007; Frenz et al., 1989; Singh and Schwarzbauer, 2012). The sequential active and passive stages, that characterize *Drosophila* VNC condensation, as reported here, have not been observed before. This complexity is probably a consequence of the specific allometric constraints that the VNC must conform to reach full functional competence (Karkali et al., 2020).

### Oscillations are an emergent property of the viscoelastic character of the VNC

The condensation of a tissue is the result of the spatially-patterned dynamics of its components (cells compression and/or neighbors exchanges), spanning micro- to millimeter scales. The directionality and magnitude of such patterning are critical for changes in the tissue fine structure and cell identities (Li et al., 2017) (Shyer et al., 2017).

We have found that during condensation the neural tissue material properties and tensional mechanics undergo progressive stereotyped changes (**Figures 2** and **3**). The embryonic VNC is very soft (as with other neural tissues (Franze et al., 2013)). However, its stiffness is neither constant, nor homogeneous. Further, we have revealed an iterated tensional pattern along the AP axis, that rises and falls following segmental structural landmarks. In short, condensation, along with the associated refinement of the VNC axonal network architecture, leads to a rigid structural configuration in equilibrium, in which tensional differences are smoothed out.

Up to now, the lack of suitable biophysical models has limited the study of the mechanics during tissue condensation. Here, we utilized a viscoelastic FE model to reproduce the experimental strain and stress maps. Given the temporal and spatial oscillatory behavior in the absence of any apparent external oscillatory signal, we resorted to a simple one-dimensional viscoelastic model of active VNC condensation. Our model not just mimics the periodic oscillations of the VNC, but it predicts the different oscillatory regimes associated to the changes of viscous and elastic mechanical properties observed during VNC condensation **(Figure 4)**. In particular, by fitting the rigidity values retrieved from the AFM measurements, our rheological model demonstrates oscillations in the absence of external oscillatory inputs. Oscillations arise as a consequence of the delayed remodeling of the tissue with respect to the compressing forces. Of course, there are multiple models that can mimic oscillatory responses, either through combining reaction-convection terms (Notbohm et al., 2016) or oscillatory polarization and alignment (Petrolli et al., 2019; Peyret et al., 2019). While we cannot discard external oscillatory inputs, we have demonstrated that the generation of oscillations can occur independently from such inputs, and that the oscillatory regime of the VNC condensation depends on its material properties and effective frictional interactions with its surroundings.

### VNC condensation requires the mechanical contribution of glia and neurons

Condensation is not symmetrical along the AP axis, progressing centripetally towards the thoracic/abdominal interphase **(Figure 7C-E)**. VNC condensation thus bears mechanical similarities to the compaction of accordion bellows, in which each pleat corresponds to a neuromere unit. Though each “pleat” in the VNC is able to contract autonomously, they are temporally and directionally coordinated across a long-range by force continuity and balance. This results in the oscillatory regimes extending throughout the VNC. We have found that this long-range continuity is created by a precise coordination of the contractile activities of neurons and glia.

In the VNC around 60 glial cells are identified per abdominal neuromere (Ito et al., 1995). Amongst them, the Subperineurial Glia (SPG), which lies beneath the outer surface of the VNC, is responsible for establishing the blood-brain barrier (BBB) (Schwabe et al., 2017). It is known that glia integrity influences VNC condensation. The ablation of hemocytes causes severe defects in SPG morphology, resulting in VNC condensation failure (Martinek et al., 2008; Olofsson and Page, 2005). Conversely, directly interfering in the activity of Rac1 or Heartless in lateral glia also blocks condensation progression (Olofsson and Page, 2005). Hypothetically, the mechanical contribution of glia to VNC condensation may be linked to their participation in casting the BBB. Indeed, we found that interfering in the contractile capability of the SPGs was sufficient to phenocopy pan-glial myosin activity depletion, both in terms of condensation **(Figure S6A)** and axon network organization (**Figure S6B**). Yet, our evaluation of the mechanical consequences of glia removal indicates that the glia does not just act as a barrier, but they also act in a manner similar to a “compression sock”, wrapping the VNC neural cortex and providing rigidity (**Figures 6** and **7**). In its absence, condensation is irregularly structured, shows a substantial reduction in its strain rates, and lacks contractile oscillations. The capability of glia to compact is strongly dependent on actomyosin contractility, and it is mainly allocated to the SPGs.

Each abdominal hemisegment of the VNC comprises around 400 neurons. Axons arrange into longitudinal connectives extending along the length of the VNC and constitute a potential force-generating source. Consistent with this proposition, the axons of longitudinal connectives loop during condensation in some metamorphic insects (Pipa, 1973) and we found that the spaces in-between neuromeres evenly subside as condensation progresses. These domains, counter-intuitively, are under tensional stress at early stages, relaxing as condensation proceeds. This might imply that the axonal network resists rather than promotes AP compaction. Indeed, ablation of neurons only marginally affects VNC condensation. However, when the contractile capability of neurons was abolished, VNC condensation failed, although strain patterns were not significantly affected. This suggests that neurons are not playing a purely passive role (**Figures 6** and **7**). Indeed, the onset of the CP2 condensation stage correlates with the initiation of synaptic neural activity (Baines and Bate, 1998). Correspondingly, mutants for Syntaxin, which lack neurotransmitter release; and the pan-neural expression of either tetanus toxin or the K+ channel Kir2.1 result in the inhibition of VNC condensation (Schulze et al., 1995).

Overall, this work reveals that the viscoelastic and biomechanical properties of the nervous system are key, in concurrence with a complex series of coordinated cellular actions, for its morphogenesis. The generation of force patterns, and the ultimate acquisition of the VNC final shape, can be assigned to the concerted actions of neurons and glia through the dynamic modulation of their cytoskeleton. The neuronal contractile capability is secondary to the glial compacting power, but necessary to direct VNC condensation along the AP axis **(Figure 7C-E)**. Finally, if we assume that VNC condensation is a way to respond to evolutionary pressure for functional optimization, we speculate that the segregation and coordination of mechanical activities between emergent neurons and glia is a key factor for natural selection.

## EXPERIMENTAL PROCEDURES

### *Drosophila* Strains and Genetics

The following stocks were used:

*w1118, fas2-GFP^CB03616^* (Dr. Christian Klämbt)

*w, elav-Gal4 [C155]* (BDSC #6923)

*w; ; repo-Gal4 / TM3, Sb[1]* (BL#7415)

*w; ; pino1::Repo-Gal4::UAS-mCD8-GFP / TM6B, Dfd-GMR-nv-YFP, Sb[1], Tb[1]* (Dr. Gerald Udolph)

*w; ; pino1::Repo-Gal4::UAS-mCD8-GFP:: His2Av-mRFP / TM6B, Dfd-GMR-nv-YFP, Sb[1], Tb[1]* (this work)

*w; ; pino1::elav-mCD8-GFP / TM6B, Dfd-GMR-nv-YFP, Sb[1], Tb[1]* (Dr. Gerald Udolph)

*w; moody-Gal4:UAS-mCD8-GFP* (Dr. Christian Klämbt)

*w; UAS-zipper RNAi / CyO* (BL#37480)

*w; His2Av-mRFP* (BL#23651)

*w; His2Av-mCherry* (Dr. Lars Hufnagel)

*w, alpha-tubulin-GFP; H2Av-mRFP* (Dr. Elena Rebollo)

*w; if / CyO; UAS-GRIM* (Dr. Todd Laverty)

In all cases, unless otherwise stated, embryos of the *w1118* strain served as controls. All crosses were performed at room temperature and after 48 hours were shifted to different temperatures as the individual experiments required.

### Sample Preparations for Immunodetection

*Drosophila* embryo dissections for generating flat preparations were performed according to (Landgraf et al., 1997). Briefly, flies maintained in apple juice-agar plates at 25°C were synchronized by repetitive changes of the juice-agar plate, with a time interval of 2 hours. All embryos laid within this time window were aged for approximately 9 hours at 29°C, or until reaching mid-stage 16 (3-part gut stage). At this point embryos were dechorionated with bleach for 1 min, poured into a mesh and rinsed extensively with water. For dissection, embryos were transferred with forceps on the surface of a small piece of double-sided tape, adhered on one of the sides of a poly-L-Lysine coated coverslip. After orienting the embryos dorsal side up and posterior end towards the center of the coverslip, the coverslip was flooded with saline (0.075 M Phosphate Buffer, pH 7.2). Using a pulled glass needle the embryos were manually de-vitellinized and dragged to the center of the coverslip, where they were attached to the coated glass with their ventral side down. An incision on the dorsal side of the embryo was performed using the glass needle from the anterior to the posterior end of the embryo. The gut was removed by mouth suction and a blowing stream of saline was used to flatten their lateral epidermis. Tissue fixation was done with 3.7 % formaldehyde in saline for 10 minutes at room temperature. After this point standard immunostaining procedures were followed.

### Immunohistochemistry

Immunostaining of flat-prepped stage 16 *Drosophila* embryos was performed using the following primary antibodies: mouse anti-Fas2 (1:100, clone 1D4, DHSB), rabbit anti-Dcp-1 (Asp216) (1:100, Cell Signaling #9578), rabbit anti-GFP tag polyclonal (1:600, Thermo Fisher Scientific) and mouse anti-Repo (1:100, clone 8D12 DHSB).

The secondary antibodies used for detection were: Goat anti-Rabbit IgG (H+L), Alexa Fluor 488 conjugate (A-11008), Goat anti-Rabbit IgG (H+L) Alexa Fluor 555 conjugate (A-21428), Goat anti-Mouse IgG (H+L) Alexa Fluor 488 conjugate (A-11001) and Goat anti-Mouse IgG (H+L) Alexa Fluor 555 conjugate (A-21422). All secondary antibodies were used in a dilution of 1:600 and were from Invitrogen.

### Confocal Image Acquisition

Flat-prepped immunostained embryos were mounted in Vectashield anti-fading medium (Vector Laboratories, USA). Image acquisition was performed on a Zeiss LSM 700 inverted confocal microscope, using a 40 X oil objective lens (1.3 NA). Z-stacks spanning the whole VNC thickness were acquired with a step size of 1 µm.

For live imaging, dechorionated stage 14 embryos were glued lateral or ventral side down on a MatTek glass bottom dish and they were covered with S100 Halocarbon Oil (Merck) to avoid desiccation. Image acquisition was performed on a Zeiss LSM 700 inverted confocal microscope, using a 25 X oil immersion lens (0.8 NA) and on a NikonA1Rsi, using a 20 X air lens (0.75 NA). Z-stacks spanning the whole VNC thickness with a 2 µm step size were acquired every 5 or 10 minutes for a total of 8-16 hours.

### Light-Sheet Imaging

Multi-view light-sheet imaging was performed on a custom-built setup. The design and imaging capabilities are similar to systems previously described (Krzic et al., 2012). Embryos were mounted in low-melting agar (0.8% w/v) filled inside FEP tube (wall thickness 0.3 mm, inner diameter 0.5 mm, refractive index 1.3) and imaged through FEP tube submerged in sample chamber filled with PBS buffer. The sample was illuminated with a light-sheet created by two long working objectives (Olympus 10 X, 0.3 NA) on the opposite side and two orthogonal fluorescence collection objectives (Nikon, water-immersion, 25 X, 1.1NA, WD 2mm). The fluorescence signal was collected and the image formed by a tube lens (Nikon, f-200 mm) on two sCMOS cameras (Hamamatsu, ORCA-Flash4.0 V2, pixel resolution 2048 × 2048, effective pixel size at object space 0.26 μm). 100 images were collected (z-resolution 1.8-2.6 μm) at 5 min time interval. Embryos were rotated 90° at each time point in order to reconstruct the full embryo morphology (see (Krzic et al., 2012) for reconstruction details).

### Image Processing

Basic confocal image processing and analyses were performed using Fiji (Schindelin et al., 2012).

Vitelline membrane autofluorescence was removed from confocal 4D images employing an ImageJ / Fiji automated macro approach (Boix-Fabres et al., 2019).

To perform the detwitching, we first generated with Matlab effectively isotropic three-dimensional reconstructions of the embryo, at each time point, using 3D linear interpolation of the z-stack obtained from imaging the ventral side of the embryo. On a single time point, we manually identified the most anterior, posterior and ventral positions of the condensing CNS. We used the built-in Matlab 3D affine transformation function to map these points to the xy-plane of the transformed image. We then chose a reference time point at the middle of the condensation process (when the VNC was positioned ventrally) and used the *imregtform* Matlab function to align the other images. This process allowed us to suppress 3D rotations due to muscle twitching. To moderate the effect of local rapid muscle contractions, we blurred the movies in space and time (Gaussian filter, pixel size 2 in x, y and time).

### Atomic Force Microscopy

Staged embryos were placed on top of positively charged glass slides to immobilize them on a rigid substrate. The embryos were immersed in PBS solution and dissected to expose the CNS allowing AFM measurements. Force-indentation curves were obtained with a custom-built AFM mounted on an inverted optical microscope (TE2000; Nikon). A 20 µm diameter polystyrene bead was glued to the end of a tip-less cantilever (nominal spring constant k= 0.01 N/m, Novascan Technologies, Ames, IA), which had previously been calibrated by thermal tune oscillation (Jorba et al., 2017). The cantilever was displaced in 3-D with nanometric resolution by means of piezo-actuators coupled to strain gauge sensors (Physik Instrumente, Karlsruhe, Germany) to measure the vertical position of the cantilever (z). The deflection of the cantilever (d) was measured with a quadrant photodiode (S4349, Hamamatsu, Japan) using the optical lever method. Before each slice measurement, the slope of a deflection-displacement d–z curve obtained from a bare region of the coverslip was used to calibrate the relationship between the photodiode signal and cantilever deflection. A linear calibration curve with a sharp contact point was taken as indicative of a clean undamaged tip. Force (F) on the cantilever was computed as Hookean linear spring:

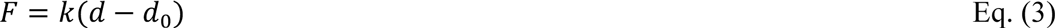

where k is the cantilever spring constant. Indentation depth δ was defined as:

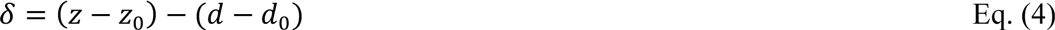

where d_0_ as is the deflection offset and z_0_ the cantilever displacement when the tip contacts the surface of the sample. F-z curves were analyzed with the Hertz contact model for a sphere indenting a semi-infinite half space:

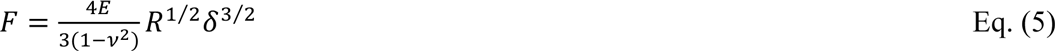

where R is the bead radius (10 µm), E is the Young’s modulus and ν is the Poisson’s ratio (assumed to be 0.5). Eq. (3) can be expressed in terms of z and d as:

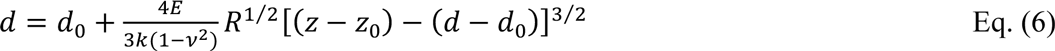

E, z_0_ and d_0_ were computed for each force–indentation curve by non-linear least-squares fitting using custom built code (Matlab). The fitting was performed for the approaching force curve up to a maximum indentation of ∼4 µm. At each measurement point, E was characterized as the average of the values computed from five force curves consecutively obtained with ramp amplitude of 20 µm and frequency of 0.3 Hz. For each embryo, the Young’s modulus (E) was measured at, at least, 3 positions along the antero-posterior axis in the midline and in lateral regions (left and right separated 20 µm from the midline).

### Laser Ablation

Laser ablation experiments were performed on a Zeiss microscope stand equipped with a spinning disk module (CSU-X1; Yokogawa), an EMCCD camera (Andor) and a custom built laser ablation system using a 355 nm pulsed laser with energy per pulse in the 20 µJoule regime and a pulse repetition of 1000 Hz (Mayer et al., 2010).

Linear ablations were performed with a 50 µm line oriented perpendicular to the VNC AP axis at different positions (intercommissural or intracommissural) between the 1^st^/4^th^ abdominal segments at different embryonic stages. The laser was focused on equally spread points (shots) on the ROI, with a density of 2 shots/µm. For each shot, 25 laser pulses were delivered. The ablation was done in a single plane, cutting the axonal network, where the entire region of interest could be acquired. To capture the rapid recoil of the ablated front, single plane images were acquired with 50 ms exposure and with a 100 ms interval between frames. Initial recoil velocity of the ablated region was computed for the estimation of mechanical stress in the tissue.

The images obtained from the laser ablations were analyzed using Fiji and Matlab. Kymographs were drawn on both sides (proximal- and distal) of the ablated line using the FIJI plugin Multi Kymograph (Schindelin et al., 2012). Both recoil velocities were calculated using a custom written routine in Matlab. The final recoil velocity (V_average_) for one ablation was computed as the average of the recoil velocities that are both proximal-oriented (V_proximal_) and distal-oriented (V_distal_):

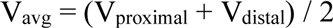

We performed an exponential fitting of experimental recoil curves *y*(*t*) to a function describing the relaxation of a viscoelastic tissue

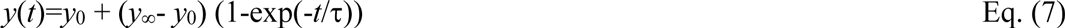

where y_0_ is the initial opening, *y*_∞_ measures the recoil increment (or tissue contractility) and τ is the characteristic time which is proportional to tissue viscosity.

### Statistical Tests

Statistical tests were performed using Matlab and estimationstats.com. Kolmogorov-Smirnov test was performed to test whether the observed values where normally distributed. When the distribution was normal, Student’s t-test was performed to estimate significance of the quantities. In case the values are not normally distributed, non-parametric Mann-Whitney U-test was performed to estimate the significance of differences between the quantities. The corresponding p-values and the method used to estimate them are mentioned in the figure legends.

### Particle Image Velocimetry (PIV)

Tissue displacements were analyzed by tracer particles, which in our experiments were EGFP-labeled Fas2 molecules and mCherry-labeled Histone2Av molecules. Flow fields were quantitatively measured using the open-source tool for Matlab PIVlab (Thielicke and Stamhuis, 2018). The software calculated the displacement of the tracers between pairs of images (sequential time points) using the Fast Fourier Transformation algorithm with multiple passes and deforming windows.

We also wrote custom software for performing the PIV analysis on the light-sheet microscopy data. This software is available upon request. From the PIV results, calculating the strain rate is then calculated by taking the spatial derivative of the PIV field after Gaussian smoothing (in both space and time) to reduce effects of noise.

### Modeling

The experimental velocity field extracted from the PIV analysis was mapped onto the closest nodes of the FE model of the VNC (**Figure 3F** and **Figure S3C**), which uses an initial geometry that resembles the VNC before condensation. Mechanical equilibrium is imposed in order to find the deformation of the whole computational domain.

A simple Maxwell rheological model was used for computing the stress values. After the FE discretization with 15800 linear hexahedral elements (**Figure S3C**), Cauchy’s equilibrium equation ∇·σ=0 yields a system of ordinary differential equations (ODEs) in terms of the nodal displacement vector u:

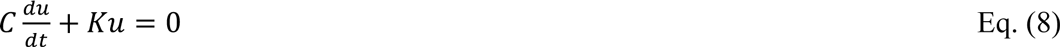

where *C* is the matrix with viscous contributions and *K* is the stiffness matrix gathering the measured elastic properties of the material. We used a Young modulus of E=75 Pa and a viscous coefficient η=500 Pa.s, which gives a similar characteristic time to the one measured through laser ablation (**Figure 3E**). The ODE system in Eq. (8) was integrated with an implicit second order accurate Crank-Nicholson scheme. The mapping of the velocities and the implementation of the FE model were done in a custom code in Matlab.

After imposition of the measured velocity field, we interpreted the resulting viscoelastic stresses of the model as the active stress field of the tissue, necessary for undergoing the condensation process. From these total stresses in the three-dimensional domain of the VNC, we extracted the compressive stress peaks along the AP axis, and fitted their evolution with a fourth inverse degree polynomial (see **Figure 3F** showing the fitted lines, and **Figure S3D** also showing the stress peaks on the kymograph). Stress peaks evolve spatially, with a magnitude that is maintained due to the VNC condensation and concomitant tissue relaxation.

Characteristic times *τ* in **Figure 3E** have been computed by fitting the gap *d*(*t*) in the recoil with an exponential function *d*(*t*) = *Ae*^-t/τ^ + *B*.

The stability analysis shown in **Figure 4** is computed from the characteristic equation of the system of delay differential equations in Eq. (1) and (2) (see (Dawi and Munoz, 2021) for further references).

## ACKNOWLEDGEMENTS

We would like to thank Christian Klämbt, Nicholas Tolwinski and Pavel Tomancak for critical reading of the manuscript, Elena Rebollo for her constant help and support at the Molecular Imaging Platform of the IBMB and Stephan Grill, MPI-CBG, for providing access to the laser ablation microscope.

KK and EMB are supported by grants BFU2017-82876-P of the Spanish Ministry of Research and Development, 2017 SGR 1199 GRC of the Generalitat de Catalunya and 2014 Fundación Ramon Areces to EMB. Work in the laboratory of TES (PT, AS and ST) was supported by a Singapore NRF Fellowship (2012NRF-NRFF001-094), an HFSP Young Investigator Grant (RGY0083/2016) and MBI Core funding. IJ and DN were supported by the Spanish Ministry of Science, Innovation and Universities (MICINN) under grant DPI2017-83721-P and the European Union’s Horizon 2020 Research and Innovation Program under the Marie Skłodowska-Curie grant 812772. JJM is financially supported by the Spanish Ministry of Science, Innovation and Universities (MICINN) and by the Generalitat de Catalunya, under the grants DPI2016-74929-R and 2017 SGR 1278, respectively.

