## Supplemental Figures for "Condensation of the *Drosophila* Nerve Cord is Oscillatory and depends on Coordinated Mechanical Interactions"

#### SUPPLEMENTAL FIGURES AND LEGENDS

### Figure S1

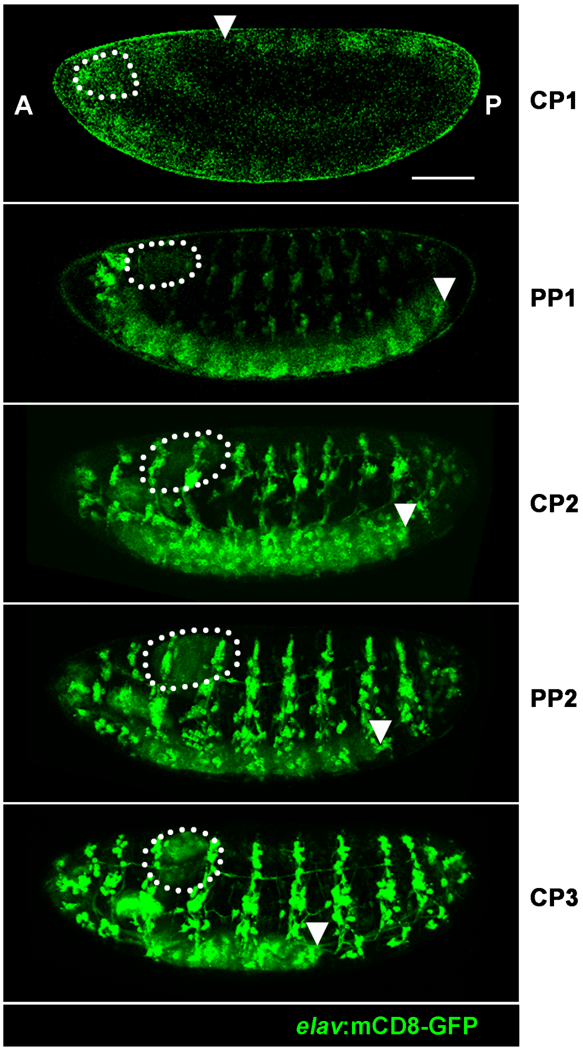


**Figure S1: VNC condensation temporal development**

Snapshots corresponding to the five phases of VNC condensation (CP1, PP1, CP2, PP2, CP3) from a time lapse **(Movie S1)** of an *elav*:mCD8-GFP embryo (lateral view) recorded by confocal microscopy. mCD8-GFP labeling marks all neural derivatives. Dotted shapes indicate the position of the brain lobes. Arrowheads denote the posterior tip of the VNC. AP axis orientation is indicated. Scale bar 50µm.

### Figure S2

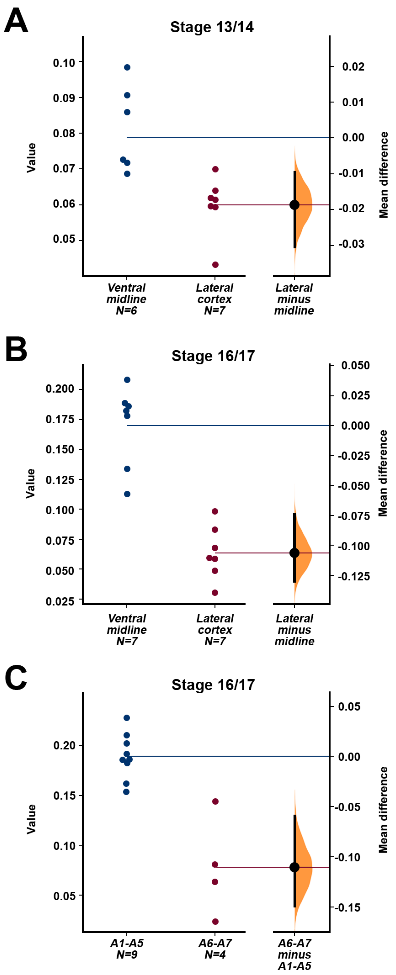


**Figure S2: Quantification of VNC material properties during condensation**

**A)** Statistical analysis of measured E (stiffness values) at different positions from stage 13-14 (estimationstats.com). Data points are shown on the left. The confidence interval is shown on right. $p < 0.05$ from Mann-Witney test. **B)** As **(A)**, but for late stages 16-17. $p < 10^{-2}$ from Mann-Witney test. **C)** As **(B)**, but comparing the E measured in anterior domains (A1-A5) with those of posterior domains (A6-A7). $p < 10^{-2}$ from Mann-Witney test.

### Figure S3

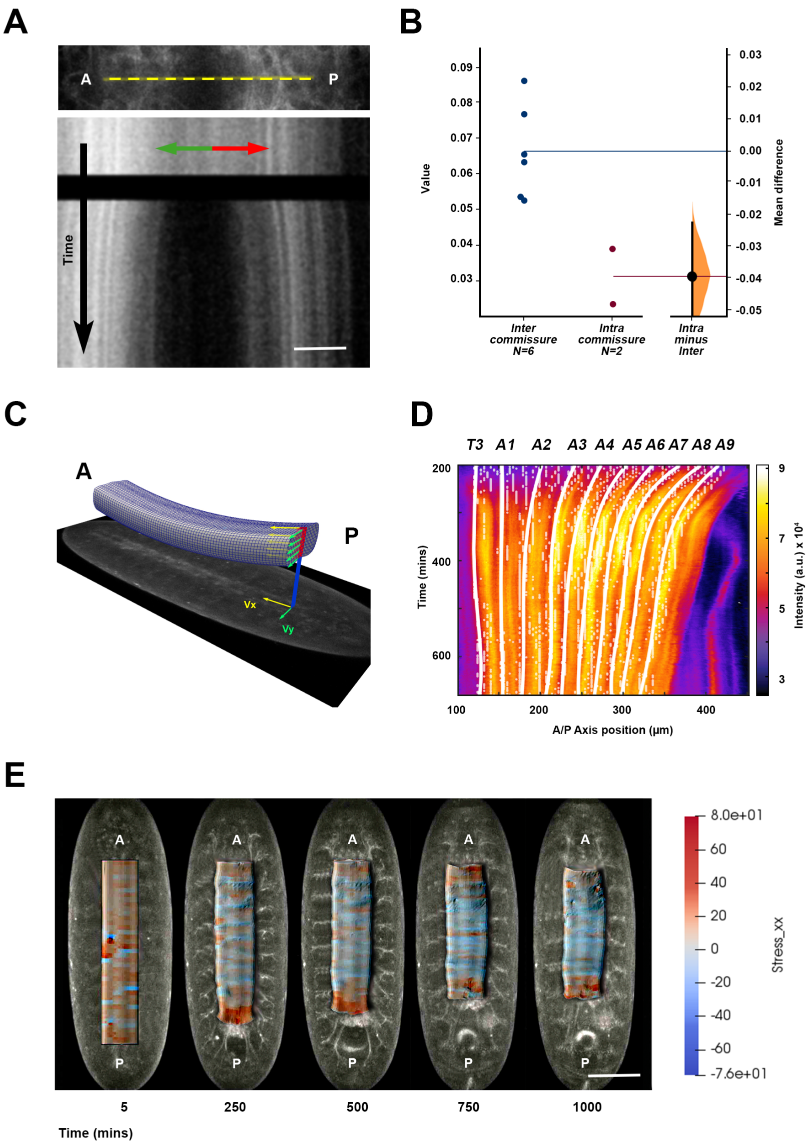


**Figure S3: Related to Figure 3. Analyses of laser microsurgery of the VNC and details of the finite elements model**

**A)** (Top) VNC tissue recoil after generating a laser cut perpendicular to the AP axis of a stage 14 embryo expressing alpha Tubulin-GFP. Yellow dashed line indicates the region of analysis. (Bottom) Kymograph of VNC recoil after laser ablation. Green (anterior) and red (posterior) arrows indicate tissue recoil directionality. The black transversal domain spans the period of laser cutting. Scale bar 10 µm. **B)** Analysis of VNC recoil speed, at inter- and intracommissural domains. Confidence interval on right. Generated using estimationstats.com. **C)** Mapping of the measured velocities from PIV onto the FE model. Each velocity on the (x, y) plane is mapped onto points of the deformed mesh with closer (x, y) positions. Nodes with non-associated velocity were deformed according to Cauchy's equilibrium equation for a viscoelastic material and discretized (see Experimental Procedures). **D)** As **Figure 3F**, but showing the points of maximum compression (minimum value of $\sigma_{xx}$) before (white dots) and after smoothing (white lines). **E)** Snapshots of deformed FE model showing contour plot of the AP normal stress $\boldsymbol{\sigma}_{xx}$ superimposed over the corresponding images (ventral view) of Fas2-GFP embryos. Scale bar 50 µm.

### Figure S4

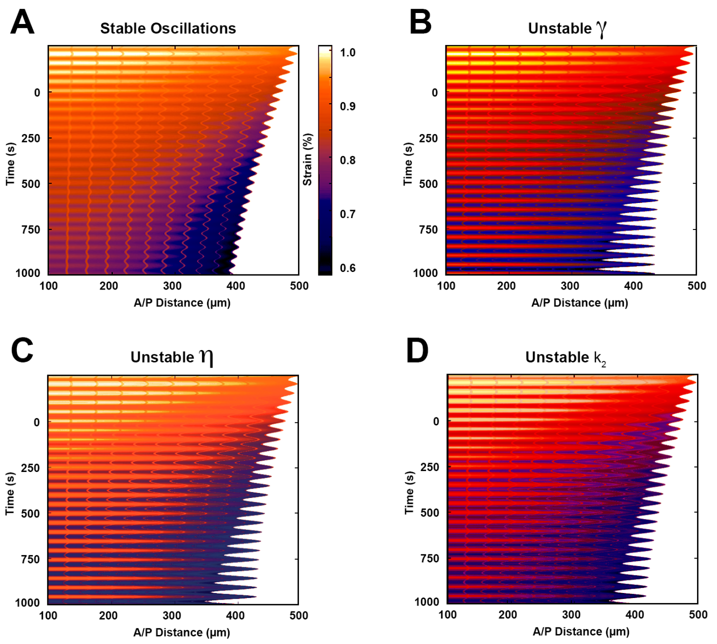


**Figure S4: Related to Figure 4. Kymograph of rest-length rheological model with delay using different material parameters and numerical simulation.**

**A)** Stable oscillation using reference values of remodeling rate $\gamma$=0.21 s$^{-1}$, viscous friction $\eta$=15 Pa.s., delay $\Delta t$ =20 s, and stiffnesses $k_1$=0.01 Pa and $k_2$= 1.9 Pa, respectively. Initial rest-length is $L_0$=0.95$l_0$,, with $l_0$ being the initial apparent length. **B)** Unstable oscillations due to increase of remodeling rate ($\gamma$=0.22 s$^{-1}$). **C)** Unstable oscillations due to increase of viscosity $\eta$=16 Pa.s. **D)** Unstable oscillations due to decrease of stiffness $k_2$= 1.8 Pa.

### Figure S5

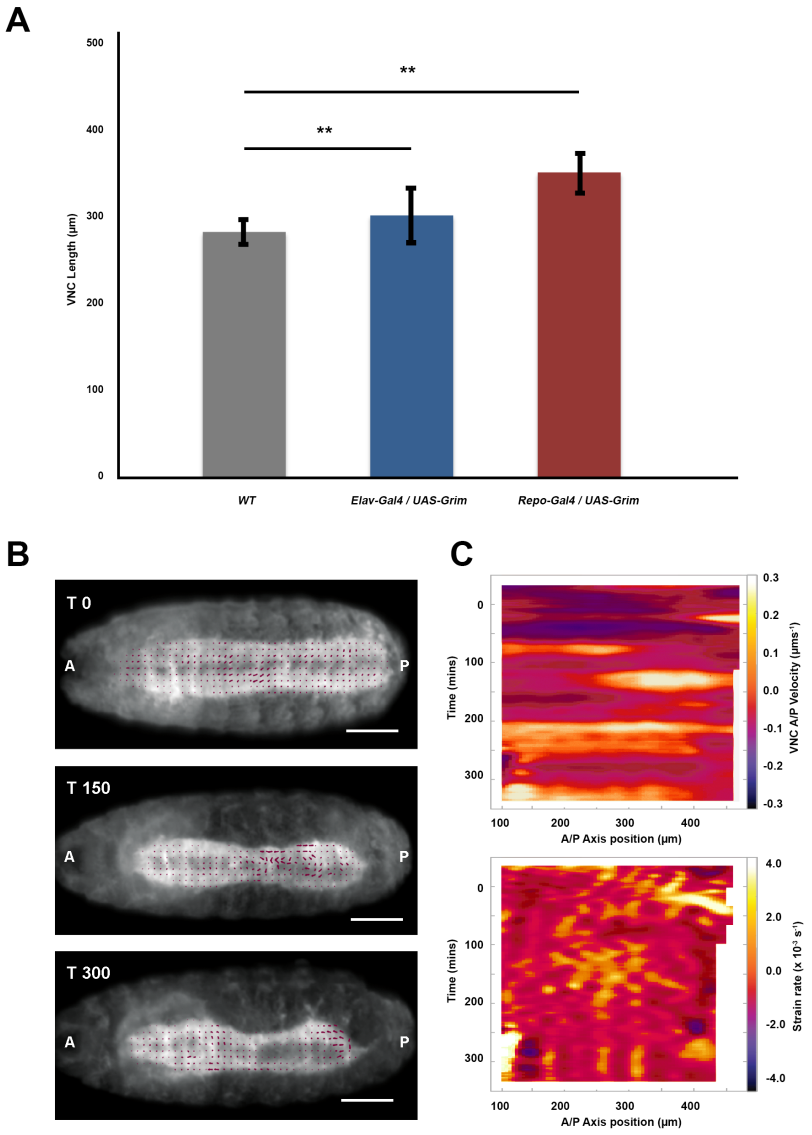


**Figure S5: Related to Figure 5. Large scale forces and local tissue dynamics are modulated by neurons and glia**

**A)** Quantification of VNC length of WT (gray), Elav-Gal4>UAS-Grim (blue) and Repo-Gal4>UAS-Grim (red) embryos, at stage 16. Bars represent mean values (n=6 embryos). $^{**}p < 10^{-2}$. **B)** Snapshots from light-sheet imaging recordings of a Repo-Gal4::UASmCD8-GFP::His2Av-mRFP>UAS-Grim embryo (ventral view) at different times of development (Stages 15-16-17) **(Movie S8)**. Magenta arrows denote local velocity trajectories from PIV analyses. AP axis orientation is indicated. Scale bar 50 μm. **C)** Velocity (as in **Figure 1C**) and strain rate (as in **Figure 2D**) kymographs for a representative Repo-Gal4>UAS-Grim embryo. No periodic oscillations were observed.

### Figure S6

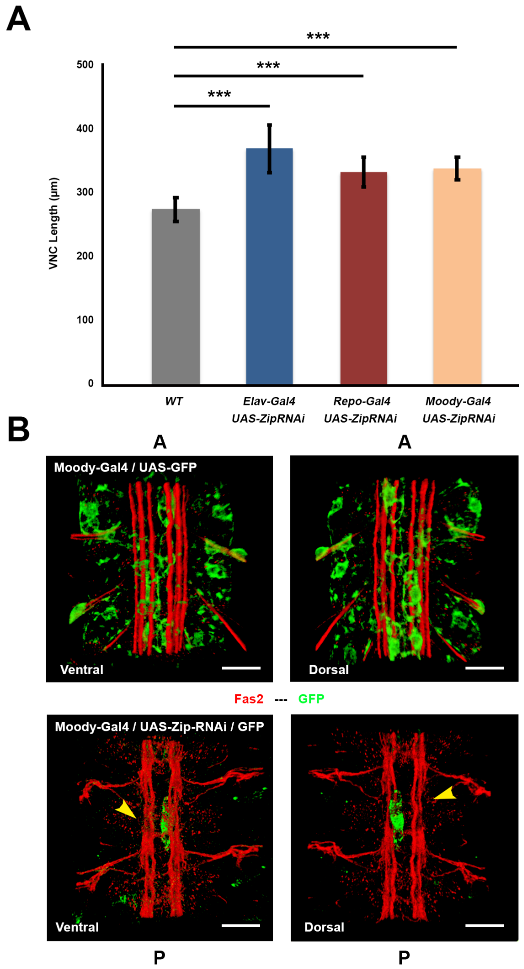


**Figure S6: Subperineural glia contractility is necessary for condensation and VNC organization.**

**A)** Quantification of VNC length of WT (gray), Elav-Gal4>UAS-Zip-RNAi (blue), Repo-Gal4>UAS-Zip-RNAi (red) and Moody-Gal4>UAS-Zip-RNAi embryos (orange), at stage 16. Bars represent mean values (n=5 embryos). ***$p < 10^{-3}$. **B)** Ventral and Dorsal 3D views of dissected, stage 16, control (top) and Moody-Gal4:UAS-mCD8-

GFP>UAS-Zip-RNAi (bottom) embryos, immunostained for Fas2 (red) and GFP (green). Yellow arrowheads point to the disrupted axonal network. AP axis orientation is indicated. Scale bar 10 μm.

#### MOVIES LEGENDS

**Movie S1. Dynamics of VNC condensation**

Time lapse of an *elav*:mCD8-GFP embryo (lateral view) recorded by confocal microscopy. mCD8-GFP labeling marks all neural derivatives. AP axis orientation is indicated. Time in hours. Scale bar 50 µm.

**Movie S2. Isotropic three-dimensional reconstructions of embryo images**

Time lapse recorded by multi-view light-sheet imaging of a live Histone2Av-mCherry embryo (ventral view). mCherry labeling marks all nuclei and was used to correct the embryo twitching. Raw data is shown on the top and “detwitched” images on the bottom. AP axis orientation is indicated. Time in hours. Scale bar 50µm.

**Movie S3. Anterior and posterior contractile oscillations (stationary domain)**

Time Lapse recording of an embryo expressing Fas2-GFP (Top - ventral view; Bottom - re-slice over the Z-axis) acquired by Confocal Microscopy. The double headed arrow points to the stationary domain where converge anterior and posterior condensation. AP axis orientation is indicated. Time in hours. Scale bar 50µm.

**Movie S4. VNC response to laser microsurgery during condensation**

Laser ablation of stage 14 embryos expressing alpha Tubulin-GFP. The recoil of intercommissural (left) and intracommissural (right) cuts are compared. Yellow lines highlight the position of the laser cuts. AP axis orientation is indicated. Time in seconds. Scale bar 20µm.

**Movie S5. VNC condensation is segmentally autonomous**

Evolution over time of a laser cut at the intercommissural space between the abdominal segments A1 and A2, of a stage 14 embryo, expressing alpha Tubulin-GFP. After ablation, the individual neuromeres (color coded dots at the bottom mark the positions of the anterior and posterior commissures of each neuromere at sequential times) continue to condense autonomously. Yellow line highlights the position of the laser cut. AP axis orientation is indicated. Time in hours. Scale bar 20μm.

**Movie S6. Three-dimensional Finite Element model (FE)**

The measured velocity field was mapped onto the FE model to reconstruct strain and stress fields. Evolution through time of contour plots of AP stresses $\sigma_{xx}$ (FE model) superimposed over experimental live images (ventral view) of an embryo expressing Fas2-GFP. AP axis orientation is indicated. Time in hours. Scale bar 50μm.

**Movie S7. Glia participate in the architectural organization of the VNC and its condensation**

Time lapse recordings of WT (Top) and Repo-Gal4>UAS-Grim (bottom) embryos in an alpha Tubulin-GFP background (ventral view) acquired by Confocal Microscopy. AP axis orientation is indicated. Time in hours. Scale bar 50μm.

**Movie S8. VNC condensation requires the mechanical contribution of glia**

Light-sheet imaging record of a Repo-Gal4::UAS-mCD8-GFP::His2Av-mRFP>UAS-Grim embryo (ventral view) at different times of development (Stages 15-17). Magenta arrows denote local velocity trajectories from PIV analyses. The VNC is significantly elongated and misshaped. AP axis orientation is indicated. Time in hours. Scale bar 50 μm.

**Movie S9. Distinct roles for neurons and glia in VNC architecture and condensation**

Time lapse recordings of embryos expressing Fas2-GFP, monitored by confocal imaging. From top to bottom, condensation dynamics in control (WT); Elav-Gal4>UAS-Zip-RNAi and Repo-Gal4>UAS-Zip-RNAi embryos. AP axis orientation is indicated. Time in hours. Scale bar 50 μm.

**Movie S10. Finite Element model of VNC condensation**

Three-dimensional representation of VNC condensation. FE model showing the evolution through time of contour plots of AP displacements (top) and experimental live images (ventral view) of an embryo expressing Fas2-GFP (bottom). AP axis orientation is indicated. Time in hours.

**Movie S11. Myosin-mediated contractility in neurons and glia is required for VNC condensation**

Finite element simulations with mapped velocities of control (WT); Elav-Gal4>UAS-Zip-RNAi and Repo-Gal4>UAS-Zip-RNAi embryos. Contour plots in the top row show the AP displacement fields and in the bottom row the elastic strains ε_xx.
